# MORPH2DIAG: Automated Structural MRI Preprocessing and Tissue Segmentation for Interpretable Machine and Deep Learning-Based Neuroanatomical Classification

**DOI:** 10.1101/2025.11.16.688711

**Authors:** Shreya C. Bangera, Libor Pospisil, Thomas Bengtsson

**Affiliations:** Department of Statistics, University of California Berkeley

## Abstract

Structural MRI provides a noninvasive window into brain morphology, yet the reproducibility and interpretability of morphometric analyses remain limited by inconsistent preprocessing, variable spatial alignment, and heterogeneous feature construction. We introduce MORPH2DIAG, a fully automated, atlas-free morphometric pipeline that integrates standardized preprocessing, tissue segmentation, spatial normalization, data-driven subtyping, and machine- and deep-learning classification within a single, modular framework. The pipeline performs intensity normalization, morphological cleanup, PCA-informed affine alignment, isotropic rescaling, and Gaussian Mixture Model (GMM) segmentation to generate quantitative gray-matter (GM), white-matter (WM), and cerebrospinal-fluid (CSF) maps. Global tissue fractions were used to derive latent neuroanatomical subtypes via unsupervised K-means clustering, revealing progressive GM–CSF gradients consistent with patterns commonly observed along normative-to-atrophic structural continua observed in neurodegeneration. To capture finer-grained spatial heterogeneity, a voxel-wise K-means parcellation yielded parcel-level intensity means and variances that served as regional morphometric descriptors. These global and parcel-level features were integrated into a unified evaluation suite comparing classical machine learning models (Random Forests, Logistic Regression, XGBoost) with lean, deep, and hybrid multilayer perceptrons (MLPs) trained using focal loss, label smoothing, stochastic weight averaging, and nested cross-validation with PCA-based dimensionality reduction. Across methods, the hybrid MLP achieved the highest macro-F1 and balanced accuracy, demonstrating strong discriminative performance for the discovered morphometric subtypes. Collectively, MORPH2DIAG establishes a fully automated, atlas-free framework that unites unsupervised structural subtype discovery with interpretable machine and deep learning, providing a reproducible foundation for MRI-based morphometric profiling and automated detection of neurodegenerative-like patterns.

## 1. Introduction

Magnetic Resonance Imaging (MRI) is a cornerstone of structural neuroimaging, providing high-resolution insight into brain morphology, tissue composition, and volumetric organization^1^. However, raw MRI data are inherently heterogeneous—affected by scanner hardware, magnetic field strength, voxel resolution, orientation, and intensity scaling—which can introduce systematic biases that confound quantitative morphometric analyses^2^. Ensuring consistency and comparability across subjects therefore requires robust preprocessing and spatial normalization as foundational steps for any downstream modeling^2^.

While established neuroimaging suites such as ANTs^3^, FSL^4^ and SPM^5^ offer comprehensive pipelines for denoising, normalization, and registration, they often entail complex dependencies, steep learning curves, and limited transparency in parameterization—factors that hinder accessibility and reproducibility. To address these challenges, we developed MORPH2DIAG, a fully automated, Python-based framework for end-to-end morphometric analysis that unifies image preprocessing, feature extraction, clustering, and classification within a single, modular workflow. The pipeline standardizes structural MRI preprocessing through intensity normalization, morphological cleanup, principal-component–based affine alignment, isotropic rescaling, and unsupervised Gaussian Mixture Model (GMM) segmentation into gray matter (GM), white matter (WM), and cerebrospinal fluid (CSF). These tissue maps yield reproducible quantitative features, global tissue fractions and voxel wise intensity distributions, that form the basis for higher-level morphometric modeling.

A key limitation in many structural MRI datasets lies in the scarcity or inconsistency of ground-truth diagnostic annotations, which often vary across studies, depend on clinical judgment, and fail to capture early or ambiguous disease states. Such label uncertainty limits the utility of purely supervised learning, as models trained on noisy or incomplete categories can overfit dataset-specific definitions rather than uncover biologically valid structure^6^. Data-driven morphometric subtyping provides an unbiased way to infer latent structural phenotypes directly from MRI features—an approach increasingly used to uncover biologically meaningful disease stages when clinical labels are unreliable or heterogeneous. These unsupervised phenotypes can then serve as stable, reproducible anchors for subsequent supervised evaluation, allowing models to test whether regional morphometric structure predicts emergent neuroanatomical organization rather than noisy clinical categories.

MORPH2DIAG adopts a hybrid learning paradigm that first discovers latent morphometric subtypes through unsupervised clustering and then evaluates their biological and statistical coherence via supervised and deep learning–based classification. In this framework, the unsupervised stage serves as a hypothesis generator—identifying intrinsic structural phenotypes based on GM–CSF trade-offs. Tissue-fraction features are clustered using K-means^7^ to reveal latent neuroanatomical subtypes reflecting progressive GM loss and CSF expansion across the cognitive continuum—from cognitively normal (CN) to mild cognitive impairment (MCI) and Alzheimer’s disease (AD)–like profiles^8^. The supervised models, in the next stage, act as hypothesis testers, assessing whether these subtypes are separable and reproducible across unseen data. The derived data-driven labels and regional parcel-wise features—representing the mean and variance of voxel intensities within each spatially defined parcel, together with global tissue fractions (GM, WM, CSF)—serve as compact morphometric fingerprints of each subject’s brain. These features preserve localized anatomical variability while reducing voxel-level noise, effectively translating high-dimensional MRI volumes into structured, biologically interpretable feature vectors suitable for downstream machine learning and deep learning models. These features are used to train predictive models spanning classical machine learning (Random Forests^9^, Logistic Regression, XGBoost^9^) and deep learning variants of multilayer perceptrons (MLPs)^10^—including lean, deep, and hybrid configurations trained with focal loss^11^, label smoothing^12^, stochastic weight averaging^13^, and nested cross-validation. The generalization performance of these classifiers thus provides an implicit validation of the underlying anatomical structure, reframing diagnosis as a mapping from emergent morphometric organization to predictive discriminability.

This integrated framework transforms unlabeled neuroanatomical data into a predictive and interpretable system for structural MRI analysis. By coupling unsupervised structure discovery with robust supervised prediction, MORPH2DIAG bridges classical morphometry and modern representation learning—offering a reproducible, scalable, and biologically grounded foundation for automated neurodiagnostics.

## 2. Data and Study Design

### 2.1 Study Design

This investigation adopted a cross-sectional design to evaluate structural brain morphology across the spectrum of cognitive aging and Alzheimer’s disease. The primary aim was to construct an automated framework capable of producing standardized, reproducible morphometric features from T1-weighted MRI data and identifying latent anatomical subtypes that reflect variation in gray-matter and ventricular structure. The analysis was organized into sequential stages that included image preprocessing, tissue-based feature derivation, clustering model comparison, and model validation. Each stage was developed to minimize manual intervention and maximize reproducibility across participants and acquisition conditions.

### 2.2 Dataset

The dataset used in this study was derived from the open-source Neurofeedback Skull-Stripped (NFBS) repository curated by the Jagust Laboratory at the University of California, Berkeley. The collection includes high-resolution T1-weighted MRI scans (1 mm³ isotropic) from adults volunteers. Crucially, NFBS does not include clinical diagnoses or cognitive labels. In this study, NFBS served as a label-free structural dataset appropriate for developing and validating MORPH2DIAG’s preprocessing, segmentation, parcellation, and subtype-discovery components. All images were distributed with precomputed intracranial masks generated through standardized skull-stripping and bias-field correction procedures. These masks remove non-brain tissue while preserving cortical and subcortical boundaries, providing anatomically consistent volumes suitable for automated morphometric analysis. All data were de-identified and obtained under institutional review board approval.

### 2.3 Structural Tissue Classes and Biological Basis

Morphometric analyses focused on three principal tissue compartments:

- **Gray matter (GM)** corresponds primarily to neuronal cell bodies and dendritic arbors. It serves as the substrate for higher cognitive processing and is the tissue most vulnerable to Alzheimer-related atrophy.
- **White matter (WM)** comprises myelinated axonal fibers that mediate long-range communication between cortical and subcortical regions. Degeneration of these tracts disrupts global network integration and contributes to cognitive decline.
- **Cerebrospinal fluid (CSF)** fills the ventricular and subarachnoid spaces and provides mechanical protection for neural tissue. Expansion of CSF volume often reflects cortical and subcortical tissue loss associated with neurodegeneration.

Their relative proportions provide a biologically interpretable summary of global structural organization, where progressive GM loss accompanied by CSF expansion forms a canonical morphometric trajectory spanning healthy aging and neurodegenerative vulnerability^1^ ^14^. Although NFBS includes no diagnostic labels, quantifying these tissue fractions enables the identification of latent morphometric continua that mirror structural patterns commonly observed across the cognitive aging spectrum.

## 3. Methods

The MORPH2DIAG framework was developed as a fully automated, end-to-end pipeline for morphometric analysis of structural MRI data, integrating image preprocessing, feature derivation, clustering, and validation in a modular sequence. Each stage was designed to preserve the biological fidelity of anatomical data while minimizing variability introduced by acquisition artifacts or operator bias. Built entirely in Python, the pipeline implements standardized intensity normalization, spatial registration, and Gaussian Mixture Model–based tissue segmentation to derive quantitative maps of gray matter, white matter, and cerebrospinal fluid. These maps serve as inputs for voxel-wise parcellation, regional feature extraction, and dimensionality reduction, followed by unsupervised clustering and supervised machine learning and deep learning–based classification. The overall workflow proceeds from data ingestion and quality control to the extraction of tissue-specific features and subsequent identification of latent structural subtypes across the cognitive continuum, establishing a transparent and extensible morphometric-to-diagnosis framework for structural neuroimaging.

### 3.1 Preprocessing

The preprocessing stage served as the foundation of the analytic workflow, transforming raw structural volumes into spatially standardized, intensity-normalized representations suitable for quantitative comparison across individuals. All images were processed in their native space to retain subject-specific geometry and avoid interpolation artifacts associated with non-linear registration. The workflow proceeded through the following major operations:

#### (1) File identification and quality assurance

All candidate anatomical volumes were recursively located within the dataset directory using pattern-based file discovery targeting standard neuroimaging formats (Neuroimaging Informatics Technology Initiative; NIfTI, *.nii* or compressed *.nii.gz*)^15^. Each file underwent automated integrity screening to verify both physical completeness and numerical validity. Checks included file-size thresholds to exclude empty or truncated images, decompression testing for corrupted archives, and validation of voxel arrays for finite numeric content. Images failing any criterion were flagged and excluded from downstream analysis. This procedure ensured that only high-fidelity, skull-stripped volumes entered the morphometric pipeline, minimizing the propagation of artifacts into derived tissue measures.

#### (2) Spatial trimming and affine preservation

To eliminate non-informative peripheral background while retaining all anatomical tissue, each brain volume was cropped to the smallest three-dimensional bounding box encompassing all non-zero voxels. The cropping operation was implemented directly on the voxel grid of the NIfTI array to preserve the original sampling resolution (1 mm³ isotropic). After trimming, the affine transformation matrix—which encodes voxel-to-world spatial mapping—was recalculated so that the physical coordinates of each voxel remained consistent with the original orientation and scaling. This step reduced computational overhead during subsequent analyses without altering any geometric relationships among voxels, thereby preserving spatial fidelity for region-based volume quantification and template alignment^3^.

#### (3) Within-brain intensity normalization

To harmonize signal intensity distributions across subjects and mitigate scanner-dependent bias, voxel intensities were standardized within brain tissue only. A binary brain mask was derived automatically from each image using percentile-based thresholding of voxel intensities followed by morphological hole-filling to ensure continuous coverage of gray- and white-matter regions. Within this mask, intensity values were normalized using a z-score transformation. This procedure preserved relative tissue contrast while equalizing global intensity scales across participants, enabling direct comparison of morphometric features derived from gray-, white-, and cerebrospinal-fluid compartments^2^.

#### (4) Optional spatial smoothing

A light isotropic Gaussian filter (σ = 0.5 voxels) was included as an optional step to suppress high-frequency noise and enhance the continuity of tissue boundaries. The Gaussian filter applies a weighted local averaging operation in which each voxel’s new intensity value is determined by its neighbors, with weights decreasing according to a Gaussian (normal) distribution centered on that voxel. The parameter σ controls the standard deviation of this distribution and thus the spatial scale of the smoothing kernel: smaller σ values (e.g., 0.5 voxels) preserve fine anatomical detail while attenuating voxel-level fluctuations, whereas larger σ values produce progressively stronger blurring and loss of structural sharpness^1^. In this study, σ = 0.5 voxels provided minimal smoothing sufficient to reduce high-frequency acquisition noise without obscuring subtle cortical thinning or ventricular expansion. To preserve native resolution for morphometric analyses, this step was disabled for all primary experiments but retained as a controlled preprocessing option for sensitivity testing.

#### (5) Output generation and provenance tracking

Each processed volume was saved in NIfTI format with an updated header and affine matrix reflecting all geometric transformations. The pipeline automatically generated comprehensive metadata logs in comma-separated value (CSV) format, documenting input and output file paths, applied operations (cropping, normalization, smoothing), and exclusion reasons. This provenance record ensures complete traceability, reproducibility, and transparency of all preprocessing decisions.^1^

Together, these operations yielded high-quality, intensity-standardized, and geometrically faithful anatomical volumes. The resulting dataset provided a consistent foundation for subsequent stages of tissue segmentation, morphometric feature extraction, and unsupervised clustering aimed at identifying structural subtypes across the cognitive aging and Alzheimer’s disease continuum.

### 3.2 Registration and Feature Extraction

After preprocessing, all skull-stripped structural images were aligned to a standardized reference template to ensure consistent orientation, scale, and spatial framing across participants.

This registration step also provided a uniform coordinate system for deriving quantitative measures of global brain geometry and signal intensity.

#### (1) Reference image selection

A multistage search routine was implemented to automatically identify an appropriate reference volume and its associated brain mask within the project repository. Priority was given to a previously generated group-average template, followed by representative scans from the Neurofeedback Skull-Stripped (NFBS) collection or, if unavailable, the standard MNI152 1 mm template. If no suitable template was found, a heuristic search located the first valid T1-weighted brain image and matching mask within the dataset. Each candidate reference was validated for file completeness, numerical integrity, and non-zero voxel content before use. This hierarchical selection strategy ensured reproducibility while maintaining anatomical correspondence with publicly available standards.^16^

#### (2) Spatial registration and affine transformation

Each preprocessed brain was aligned to the chosen reference space using a principal-component and center-of-mass (PCA-COM) alignment procedure^3^. A provisional brain mask was first derived from the intensity histogram of each volume to isolate intracranial tissue. Voxel coordinates within this mask were converted to physical (world-space) coordinates using the image’s affine transformation matrix. Principal component analysis of these coordinates defined the dominant anatomical axes of the subject brain, which were then matched to the reference axes by computing a rigid rotation matrix (R) and translation (T) that minimized orientation and positional mismatch.

An optional isotropic scale factor, s = (V_ref_ / V _src_) ^⅓^ was applied to harmonize global brain size, where V_src_ and V_ref_ represent the intracranial volumes of the source and reference brains, respectively. The final 4 × 4 affine transform W=T_2_RST_1_ combined rotation, translation, and scale, preserving correct voxel-to-world correspondence^3^.

#### (3) Resampling and spatial standardization

The transformed images were resampled onto the reference grid using trilinear interpolation for anatomical volumes and nearest-neighbor interpolation for binary masks, thereby preserving discrete tissue boundaries. All registered outputs were written in NIfTI format with updated affine and header metadata that accurately described their position in template space. This process produced uniformly oriented, voxel-aligned volumes suitable for direct morphometric comparison across individuals.

#### (4) Quantitative feature extraction

Global morphometric and intensity-based features were automatically computed from each registered image:

- **Brain volume (mm³):** number of intracranial voxels multiplied by voxel volume.
- **Voxel count and bounding-box dimensions:** spatial extent of the brain region along each anatomical axis.
- **Center of mass:** mean voxel coordinate of the intracranial mask in template space.
- **Mean and standard deviation of intensity:** summary measures of tissue-signal distribution within the brain mask.
- **Percentile statistics (1st–99th):** robust indicators of global intensity range minimizing the influence of extreme values.

These measures provided concise descriptors of whole-brain morphology and intensity normalization stability, forming the quantitative basis for subsequent clustering analyses^2^.

### 3.3 Tissue Segmentation and Volumetric Quantification

After registration to the common reference space, each intensity-normalized anatomical volume was segmented into the principal tissue compartments—cerebrospinal fluid (CSF), gray matter (GM), and white matter (WM)—to obtain biologically interpretable, tissue-specific volumes and probability maps. The segmentation framework combined adaptive intracranial masking, probabilistic intensity modeling, and morphological refinement to ensure robustness across heterogeneous image contrasts and residual noise.

#### (1) Intracranial mask generation

A coarse brain mask was first generated directly from the registered T1-weighted image using intensity percentile thresholding. Voxels above the 5th-percentile of the finite intensity distribution were identified as likely brain tissue, and the preliminary mask was refined by sequential hole-filling, morphological opening, and closing operations to remove small discontinuities and fill sulcal gaps. This adaptive procedure excluded background air and non-brain structures while preserving contiguous cortical and subcortical regions. The resulting mask defined the domain for tissue modeling and ensured that mixture fitting operated exclusively within intracranial voxels.

#### (2) Intensity standardization and trimming

Within the intracranial mask, voxel intensities were extracted and clipped to the 0.5th–99.5th percentile range to minimize the influence of extreme tails and residual artifacts (e.g., noise spikes or partial-volume voxels at tissue boundaries). This trimming stabilized subsequent parameter estimation without altering the relative separation between tissue classes^2^.

#### (3) Probabilistic modeling with a three-component Gaussian Mixture Model

Tissue classification was achieved by fitting a three-component Gaussian Mixture Model (GMM) to the one-dimensional distribution of in-brain intensities. The model represents the empirical distribution as a weighted superposition of three normal densities:

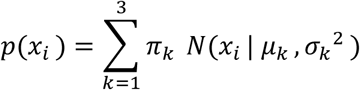

where x_i_ is the intensity of voxel *i*, π_k_ are non-negative mixture weights that sum to 1, and μ_k_, σ ^2^ are the mean and variance of class *k*. Parameters were optimized via the Expectation–Maximization (EM) algorithm^1^, alternating between:

- an *E-step* computing voxel-wise posterior probabilities p(k∣x_i_) given the current parameters, and
- an *M-step* updating π_k_, μ_k_, σ ^2^ to maximize the log-likelihood of the observed data.

Initialization by intensity quantiles provided stable convergence, and variance floors prevented degenerate solutions. After convergence, components were ordered by ascending mean intensity and assigned to CSF (low), GM (intermediate), and WM (high), consistent with T1-weighted contrast hierarchy.

#### (4) Posterior probability mapping and hard labeling

For every intracranial voxel, posterior probabilities were recomputed using the final model parameters, producing three continuous probability maps (CSF, GM, WM) whose values sum to 1. A discrete label map was then obtained by maximum-posterior classification within the mask (background = 0). This dual output preserves both uncertainty information and categorical boundaries: soft maps are valuable for partial-volume assessment, whereas hard labels support direct volumetric computation^1^.

#### (5) Volumetric quantification

Tissue volumes were calculated from the hard-label image as V_t_ = v_voxel_ × N_t_, where N_t_ is the number of voxels labeled as tissue *t* and v_voxel_ is the physical voxel volume derived from the image affine (absolute determinant of its 3 × 3 spatial sub-matrix). Volumes were reported in mm³ as CSF_mm³_, GM_mm³_, and WM_mm³_, forming subject-level quantitative summaries of intracranial composition.

#### (6) Biological Interpretability

The Gaussian Mixture Model–based segmentation framework reflects the intrinsic tri-modality of T1-weighted magnetic resonance imaging, which differentiates tissue classes by their underlying biophysical composition. In this representation, voxels characterized by low-intensity values correspond predominantly to cerebrospinal fluid (CSF), a compartment rich in water and devoid of cellular structure, while intermediate intensities represent gray matter (GM), composed mainly of neuronal cell bodies, dendrites, and synaptic processes. High-intensity voxels correspond to white matter (WM), reflecting the greater lipid content and myelination of axonal fibers^1^. By probabilistically modeling this tri-modal intensity distribution, the approach captures the essential physiological contrasts that define the human brain’s macroscopic architecture.

Across participants, systematic variation in the relative volumes of these three compartments carries well-established biological meaning. Reductions in GM and WM volumes, when accompanied by corresponding increases in CSF, provide a quantitative signature of neurodegenerative tissue loss, as seen in aging and Alzheimer’s disease. GM atrophy reflects neuronal and dendritic degeneration, particularly in association cortices and hippocampal regions, whereas WM reduction signals demyelination and axonal disconnection that compromise network integration. The expansion of CSF within the ventricular and subarachnoid spaces occurs as a passive response to this parenchymal loss, serving as a macroscopic indicator of disease progression. Thus, the derived tissue volumes from this segmentation pipeline offer not only a geometric description of brain composition but also a biologically interpretable measure of structural integrity and pathological change^17^.

### 3.4 Model Development and Clustering Framework

Following tissue segmentation, the resulting cerebrospinal fluid (CSF), gray matter (GM), and white matter (WM) volumes were used to derive population-level morphometric phenotypes through unsupervised clustering. The objective of this stage was to identify latent structural subtypes that capture inter-individual variability in global brain composition across the cognitive spectrum. The analytical workflow consisted of three stages: feature normalization, model fitting across multiple cluster cardinalities, and quantitative model selection based on complementary goodness-of-fit criteria.

#### (1) Feature construction and normalization

Each participant’s total intracranial volume was first obtained as the sum of GM, WM, and CSF compartments. To account for head-size variability and scanner-dependent scaling, all analyses were performed using fractional tissue volumes—that is, each tissue volume divided by the total intracranial volume. These normalized fractions (GM fraction, WM fraction, CSF fraction) formed a three-dimensional morphometric signature describing the relative composition of neural and non-neural compartments. Prior to clustering, features were standardized to zero mean and unit variance to ensure equal weighting across dimensions and to prevent any one compartment from dominating the solution due to differences in numerical range.

#### (2) Clustering algorithms

Two complementary unsupervised learning algorithms were evaluated across a range of candidate cluster numbers (k = 2 to 5).

1. **K-Means clustering** was used as a baseline partitioning method, grouping individuals into discrete, non-overlapping clusters based on proximity in the standardized feature space. The algorithm iteratively reassigns points to minimize the total within-cluster variance, with multiple random initializations to avoid local minima. Cluster quality was assessed using the silhouette coefficient, which quantifies how well each subject matches its own cluster compared with neighboring ones.
2. **Gaussian Mixture Modeling (GMM)** was used to provide a probabilistic alternative that accommodates overlapping clusters and unequal covariance structures. Instead of imposing hard boundaries, GMM assigns each subject a probability of belonging to each cluster, allowing for uncertainty near transitions. Model complexity was automatically penalized through the Bayesian Information Criterion (BIC), which favors parsimonious solutions that balance fit accuracy with model simplicity.

#### (3) Model selection and evaluation

All candidate models were fit across the same cluster range, and their performance metrics were compiled into a comparative score table. Three selection strategies were implemented.

1. First, a **silhouette-based approach** prioritized compact, well-separated K-Means clusters with the highest silhouette score.
2. Second, a **BIC-based approach** selected the GMM configuration with the lowest information criterion value, favoring models that best explained the data while avoiding overfitting.
3. Finally, a **hybrid strategy** combined both criteria: GMM solutions were preferred only when their BIC improvement over other models was substantial (≥ 25 points) and not accompanied by a large decline in silhouette quality (≤ 0.02). This multi-metric evaluation ensured that the chosen solution was both statistically robust and geometrically coherent.

All model comparisons were restricted to valid fits producing at least two non-degenerate clusters, and the winning configuration was retained for downstream analyses

#### (4) Cluster labeling and biological interpretability

To align the computational output with neurobiological meaning, clusters were reordered by increasing mean CSF fraction. This ordering places subjects with higher gray- and white-matter preservation (low CSF) toward one end of the continuum and those with enlarged CSF compartments (reflecting tissue loss and ventricular expansion) toward the other^17^. For Gaussian Mixture Models, each subject also retained a cluster-membership probability, representing confidence in its assignment. This probabilistic labeling captures transitional morphologies and supports interpretation of progressive or mixed structural phenotypes.

#### (5) Outcome

The final clustering framework provided a scalable and interpretable representation of individual differences in global brain composition. By jointly modeling GM, WM, and CSF fractions, it delineated structural subtypes that potentially correspond to normative, atrophic, and transitional anatomical profiles along the continuum of cognitive aging and Alzheimer’s disease. The integration of both K-Means and GMM methodologies ensured sensitivity to discrete and continuous variation alike, yielding a stable, biologically coherent basis for the validation and statistical comparisons presented in the next section.

### 3.5 Model Validation and Supervised Classification

The final analytical stage aimed to evaluate the biological and predictive validity of the data-driven morphometric clusters obtained from the unsupervised modeling framework. This involved translating cluster assignments into interpretable diagnostic categories, generating regional features through spatial parcellation, and benchmarking multiple supervised learning models to assess how well global and regional morphometric patterns could discriminate between clinical strata along the continuum of cognitive aging and Alzheimer’s disease.

#### (1) Cluster relabeling and diagnostic mapping

Numeric cluster identifiers were transformed into interpretable diagnostic categories by ranking clusters according to their mean cerebrospinal-fluid (CSF) fraction, a morphometric indicator of parenchymal loss. Higher mean CSF fractions were taken to represent greater degrees of neurodegeneration, consistent with ventricular and sulcal expansion accompanying cortical and subcortical atrophy^17^. This ordered continuum of tissue integrity was mapped to diagnostic labels in descending order of structural preservation: **Cognitively Normal (CN) → Mild Cognitive Impairment (MCI) → AD/MCI (intermediate phenotype) → Alzheimer’s Disease (AD)**.

When clustering produced more than four morphometric groups, additional categories were appended to maintain the ordinal progression of severity while preserving distinct anatomical signatures. These biologically grounded label assignments established a consistent interpretive framework linking data-driven morphometric patterns to clinically recognizable disease stages and provided the reference classes for subsequent supervised modeling of structural degeneration patterns.

#### (2) Data-driven spatial parcellation

To extract regional structural features without dependence on predefined atlases, a data-driven parcellation was created directly from the intensity structure of the preprocessed images. A representative T1-weighted brain volume was selected as a reference template. Non-brain voxels were removed by intensity thresholding, retaining the top 95 % of intracranial voxel intensities to produce a binary mask encompassing cortical and subcortical gray- and white-matter regions.

The three-dimensional voxel coordinates within this mask were then clustered using the K-Means algorithm to produce approximately 100 spatially compact parcels of roughly uniform volume. This unsupervised partitioning creates anatomically agnostic yet topologically continuous regions that adapt to the intrinsic geometry of the dataset rather than conforming to externally imposed atlas boundaries.

#### (3) Regional feature extraction and aggregation

Each subject’s registered anatomical image was resampled to the parcellation grid to ensure affine and voxel-wise alignment. Within each parcel, two descriptive statistics were computed: the mean intensity, representing the central tissue contrast of the region, and the intensity variance, reflecting local heterogeneity and boundary sharpness. These metrics capture complementary information—mean intensity relates to overall tissue composition and myelination, while variance provides a proxy for microstructural uniformity and partial-volume effects^16^.

For robustness, parcels containing no valid voxels were retained with zero-filled entries to maintain fixed feature dimensionality. To integrate global context, the normalized GM, WM, and CSF fractions (previously derived from the tissue-segmentation stage) were appended to the feature matrix. The resulting feature set thus combined regional parcel-wise descriptors and global tissue metrics, jointly characterizing both localized and system-wide anatomical variability. Columns corresponding to parcels that were consistently zero across participants were removed to minimize noise and redundancy. The final feature table, including subject identifiers and diagnostic labels, was archived as a reproducible dataset for subsequent modeling.

#### (4) Dimensionality reduction via principal component analysis

Given the high degree of correlation and potential collinearity among parcel-wise intensity features, Principal Component Analysis (PCA) was employed to transform the feature space into a low-dimensional, orthogonal basis that captures the dominant axes of morphological variance across subjects. All features were standardized to zero mean and unit variance before decomposition to ensure comparable scaling across measures.

The PCA model retained either the first 20 components or the number required to explain at least 95 % of the total variance—whichever criterion yielded fewer dimensions—thus preserving the majority of meaningful structural information while discarding redundant or noise-driven variance. The resulting principal components provided a compact representation of large-scale anatomical gradients, such as patterns of cortical thinning, ventricular enlargement, and global gray–white matter contrast shifts. This transformation enhanced the interpretability and computational efficiency of subsequent classification analyses by emphasizing coherent morphological trends and suppressing parcel-specific fluctuations.

#### (5) Supervised learning and cross-validated model comparison

To assess the discriminative power of the derived morphometric features, three independent supervised classification models were trained and evaluated using stratified five-fold cross-validation to preserve class balance across folds:

- **Random Forests**: an ensemble of 500 decision trees using balanced class weighting to mitigate group-size imbalances and automatic out-of-bag regularization for stability;^9^
- **Regularized Logistic Regression (LRCV)**: a penalized linear classifier with cross-validated optimization of the regularization strength (*C*), implemented with balanced class weights and up to 2000 training iterations; and
- **Extreme Gradient Boosting (XGBoost)**: a gradient-based ensemble model configured with moderate depth (*max_depth = 4*), conservative learning rate (*η = 0.05*), and stochastic subsampling (*subsample = 0.8*) to prevent overfitting while capturing nonlinear interactions.^9^

Each model was embedded in a standardized preprocessing pipeline including feature scaling to ensure consistency of evaluation. Cross-validation was repeated across five folds, and classification accuracy was averaged to provide an unbiased estimate of generalization performance.

#### (6) Final model training and diagnostic performance evaluation

The model with the highest mean cross-validated accuracy was selected as the final predictive framework. It was retrained on 80 % of the dataset and evaluated on the remaining 20 % held-out test set using stratified sampling to preserve class proportions. Prediction performance was summarized through standard metrics including class-wise precision, recall, and F1-score, averaged over all diagnostic categories.

The combined pipeline—spanning cluster relabeling, adaptive parcellation, regional feature extraction, dimensionality reduction, and supervised model benchmarking—provides an integrated validation framework linking unsupervised structural subtypes to clinically interpretable diagnostic categories.

### 3.6 Deep-Learning Classifiers

To complement the supervised machine-learning analyses described in Section 3.5, a suite of deep-learning models was implemented to test whether nonlinear feed-forward architectures could capture higher-order dependencies between regional morphometric features and diagnostic status. All networks operated on the same leakage-controlled feature space derived from parcel-wise mean and variance statistics combined with global gray-matter (GM), white-matter (WM), and cerebrospinal-fluid (CSF) fractions. This unified framework enabled a controlled comparison of architectural depth, regularization, and representational capacity across three model families—Lean, Deep, and Hybrid MLPs—to evaluate how hierarchical nonlinear transformations improve discrimination along the Alzheimer’s disease continuum.

#### (1) Input Representation and Latent Embedding

Prior to network training, all features—including parcel-wise mean and variance statistics combined with global GM, WM, and CSF fractions—were standardized to zero mean and unit variance within each training fold to ensure consistent scaling across subjects. Dimensionality reduction was performed exclusively within the cross-validation loop to prevent information leakage. A Principal Component Analysis (PCA) embedding was used as the sole latent representation method. The number of components was dynamically selected from a grid of candidate dimensions (e.g., 32, 64, 128), retaining the most informative morphological gradients while minimizing redundancy. Each PCA model was fit only on the training data of a given fold and then applied to the corresponding validation subset, ensuring strict separation between training and evaluation data during model optimization.

#### (2) Network architectures

Three multilayer perceptron (MLP) architectures were implemented to systematically modulate representational depth and regularization while preserving a consistent computational backbone.^18^ Each network followed the standardized internal block structure:

**LayerNorm → Linear → GELU → Dropout, repeated per hidden layer, followed by a final Linear → Softmax projection.**

- Lean MLP — A lightweight two-layer configuration (e.g., 128 → 64 → C units) emphasizing stability and generalization under limited parameter count. Its compact architecture minimizes variance and serves as a regularized baseline.
- Deep MLP — A high-capacity model with three hidden layers (e.g., 256 → 256 → 128 → C units) designed to capture higher-order nonlinear dependencies between morphometric features after PCA compression. Additional depth increases flexibility but requires stronger regularization to prevent overfitting.
- Hybrid MLP — An intermediate configuration (e.g., 256 → 128 → C units) combining the broader representational width of the Deep MLP with the shallow depth and stronger dropout regularization of the Lean MLP. This design balances model expressivity and generalization.

All networks employed the Gaussian Error Linear Unit (GELU) activation for smooth nonlinear transformation, Layer Normalization at the input to stabilize gradient flow across features, and a final linear classification head with a softmax output of size C (4 diagnostic classes).

#### (3) Regularization and optimization strategy

A unified training protocol was designed to ensure stability, fairness across architectures, and robustness against class imbalance:

- **Loss Function** — Primary objective: *label-smoothed focal cross-entropy*, combining focal weighting (γ = 1.5) with label smoothing (ε = 0.10) to emphasize harder samples while preventing over-confidence.
- **Class Weighting** — Per-fold inverse-frequency class weights applied within the loss function to counteract under-represented diagnostic categories.
- **Feature-space MixUp** — Convex interpolation between paired samples (α ≈ 0.2) introduced stochastic linearity in latent space and reduced overfitting by augmenting the training manifold.
- **Dropout Regularization** — Dropout (0.2 – 0.35) applied after every hidden layer encouraged sparse activations and reduced co-adaptation among neurons.
- **Optimizer and Learning Schedule** — All networks used **AdamW** (decoupled weight decay = 1×10⁻⁴ – 5×10⁻⁴) with a **One-Cycle learning-rate** schedule (peak LR ≈ 2–3×10⁻³), ensuring fast convergence followed by gradual annealing.
- **Gradient Stability** — Gradients were clipped at ‖g‖₂ ≤ 5 to suppress exploding updates in deeper models.
- **Early Stopping** — Monitored macro-F1 on the validation set; training terminated after ∼15–20 epochs without improvement.
- **Stochastic Weight Averaging (SWA)** — After epoch 140, moving-average weights were maintained to flatten the loss landscape; the checkpoint achieving maximal validation macro-F1 was retained for evaluation.

This consistent optimization framework ensured that improvements in performance were attributable to architectural differences rather than training variance.

#### (4) Hyperparameter selection

All hyperparameters were optimized using a nested cross-validation design to prevent data leakage and overfitting.

Within each outer training fold, an inner grid search tuned:

| <u>Parameter</u> | <u>Search Space</u> | <u>Purpose</u> |
| --- | --- | --- |
| PCA components | {32, 64, 128} | Dimensionality of latent morphometric embedding |
| Hidden-layer width | {128, 256} (Lean/Hybrid); {256, 512} (Deep) | Model capacity |
| Dropout rate | {0.20, 0.35} | Regularization strength |
| Weight decay | $\{1 \times 10^{-4}, 5 \times 10^{-4}\}$ | Parameter shrinkage |

The inner-fold macro-F1 score served as the selection criterion. All preprocessing (standardization + PCA) was re-fit exclusively on training data within each fold, and the optimal configuration was re-trained on the full fold’s training set before outer-fold evaluation.

#### (5) Cross-validation design and evaluation metrics

A five-fold stratified outer cross-validation procedure was employed to estimate generalization accuracy. Within each fold, the network was trained on 80 % of subjects and validated on the remaining 20 % using the fold-specific hyperparameters described above. Performance was summarized by macro-averaged F1 as the principal measure of balanced classification, complemented by accuracy and balanced accuracy.

Per-class precision, recall, and F1 were computed per fold, and confusion matrices were aggregated across folds to visualize systematic confusions between cognitive strata.

## 4. Results

### 4.1 NFBS Dataset: Structural Overview

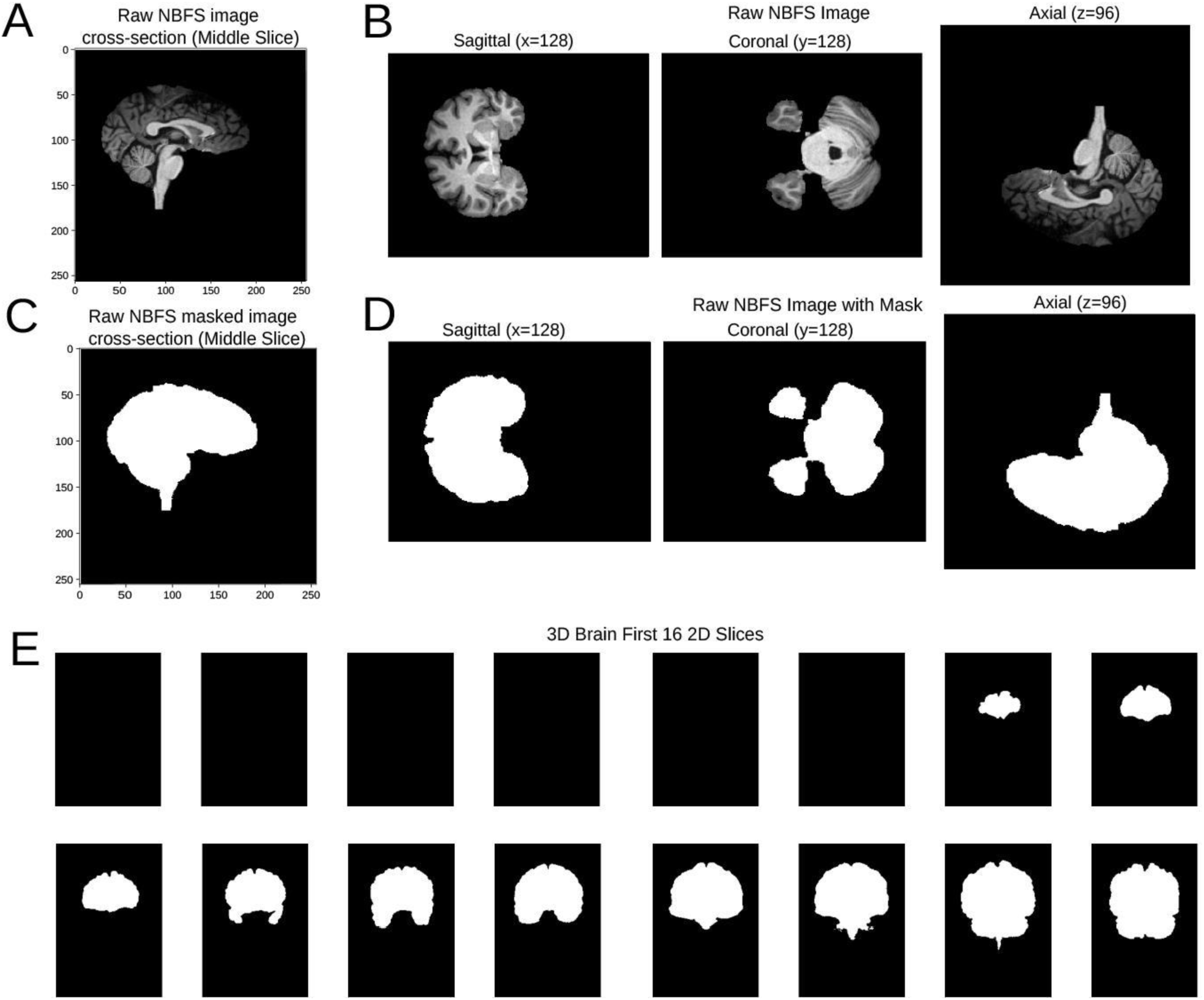

To validate and illustrate the preprocessing pipeline, we first examined representative cross-sectional views (Fig. 1A) from the Neurofeedback Skull-Stripped (NFBS) dataset. Fig. 1B depicts high-resolution axial, coronal, and sagittal slices of a single NFBS T1-weighted brain volume, demonstrating the preserved anatomical detail following intensity normalization and morphological cleanup. For comparison, the corresponding binary brain mask (Fig. 1C,D) is displayed in identical orientations, confirming accurate skull-stripping and clear delineation of cortical and subcortical boundaries. To provide a volumetric perspective, the first 16 consecutive axial slices of the same three-dimensional image are also shown (Fig. 1E). This series captures the progressive anatomical transition from inferior to superior cortical regions and highlights the spatial continuity of the normalized signal intensity across the entire brain volume. The corresponding mask slices confirm consistent foreground segmentation and the absence of residual extracerebral artifacts.

### 4.2 Preprocessed NFBS Images and Intensity–Volume Distributions

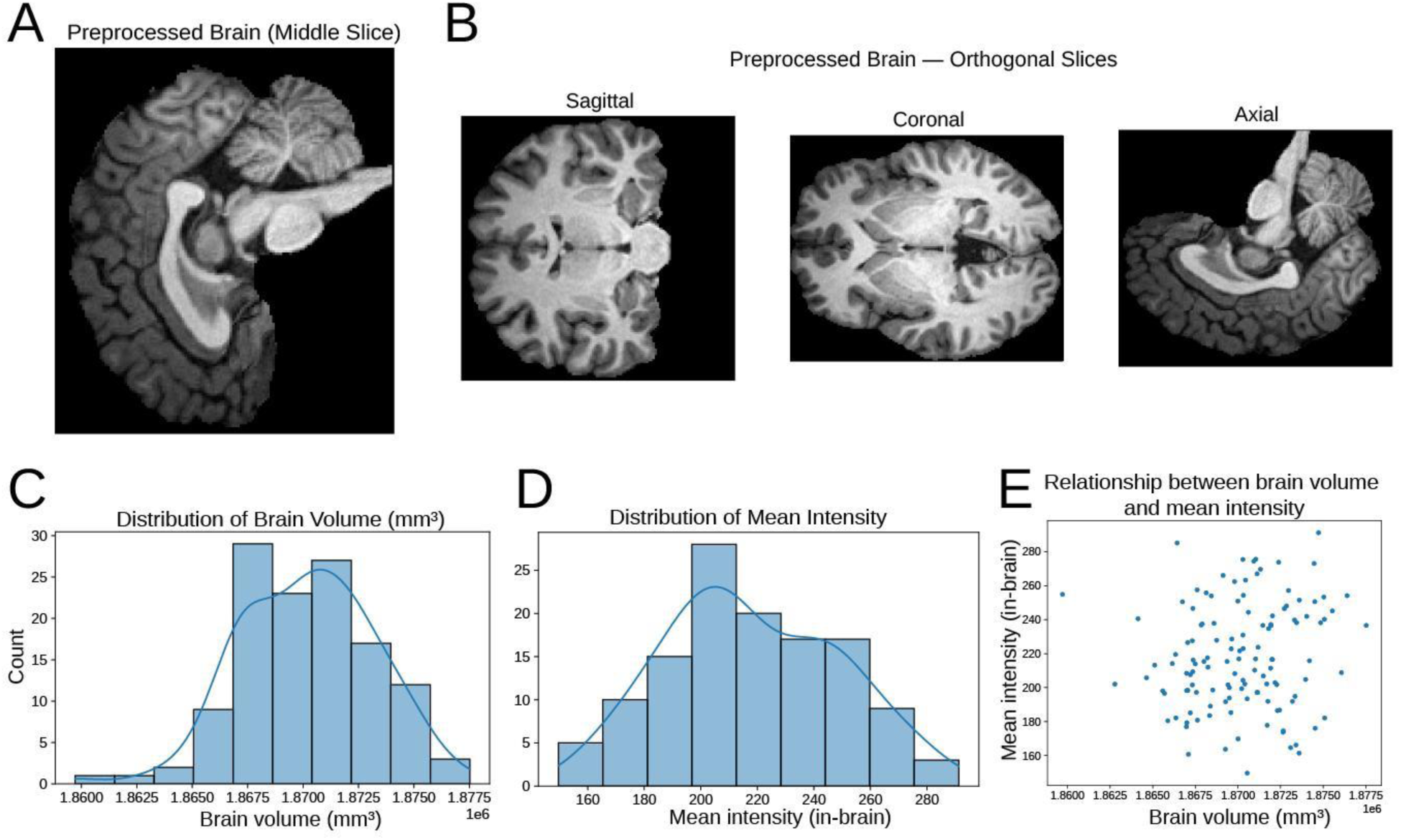

Fig. 2A shows the same axial NFBS slice as in Fig. 1A after preprocessing, illustrating the effect of intensity normalization, morphological cleanup, and affine alignment on structural clarity. Gray–white matter boundaries appear sharper, non-brain tissue is effectively removed, and overall contrast homogeneity is improved relative to the raw image. Fig. 2B presents the corresponding axial, coronal, and sagittal planes following full pipeline implementation, confirming that the preprocessing preserves fine-grained cortical and subcortical morphology while minimizing residual intensity artifacts across orientations. We summarize the global image statistics: the brain volume distribution (Fig. 2C) approximates a normal curve across participants, the mean intensity histogram (Fig. 2D) also follows an approximately Gaussian profile after normalization, and the scatterplot of mean intensity versus brain volume (Fig. 2E) reveals a clear positive correlation—indicating that subjects with larger intracranial volumes tend to exhibit proportionally higher normalized signal means. Collectively, these results validate the consistency and robustness of the preprocessing pipeline in generating standardized, quantitatively comparable MRI inputs for subsequent modeling.

### 4.3 Tissue Segmentation and Feature Extraction

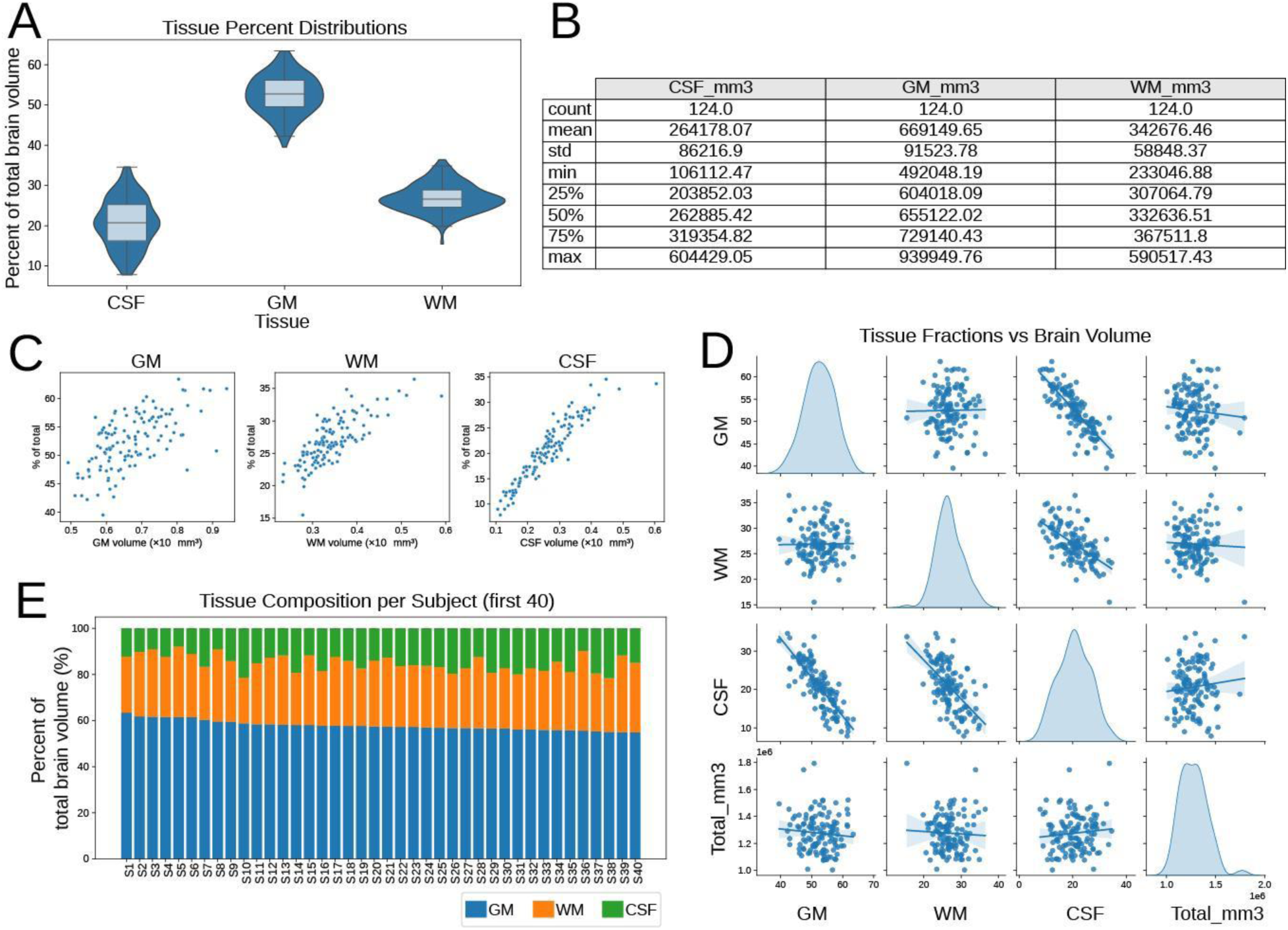

Each preprocessed MRI volume was segmented into gray matter (GM), white matter (WM), and cerebrospinal fluid (CSF) compartments using Gaussian mixture–based classification to derive subject-level morphometric features. The resulting tissue percentage distributions showed a biologically coherent hierarchy—GM occupying the largest proportion of total brain volume, followed by WM and then CSF (Fig. 3A). Summary statistics confirmed this pattern, with mean GM ≈ 669 k mm³, WM ≈ 343 k mm³, and CSF ≈ 264 k mm³ across 124 subjects, indicating realistic volumetric ratios and moderate inter-individual variability (Fig. 3B). When absolute tissue volumes were plotted against their percentage contributions (Fig. 3C), all three tissue types exhibited strong positive trends, confirming proportional scaling after normalization. However, the scatter tightness systematically increased from GM → WM → CSF. This pattern reflects underlying biological variability: GM shows the greatest inter-subject dispersion due to differences in cortical thickness and folding; WM exhibits intermediate variability related to tract density and myelination; and CSF displays the most compact relationship because its intracranial compartment is relatively small and scales almost linearly with head size. The progressively tighter scatter thus captures genuine anatomical scaling rather than preprocessing artifacts. Pairwise comparisons of tissue volumes (Fig. 3D) further highlight these relationships. GM and WM volumes were positively correlated, consistent with coordinated growth and atrophy patterns across parenchymal tissues. In contrast, CSF volumes were negatively correlated with both GM and WM, reflecting the compensatory expansion of ventricular and subarachnoid spaces that accompanies reductions in parenchymal tissue. Finally, subject-level stacked bar plots illustrate individual differences in tissue composition (Fig. 3E), showing that despite modest variability, GM consistently dominates total brain volume. Together, these findings confirm that the segmentation and feature-extraction pipeline produced anatomically interpretable, quantitatively reliable tissue measures suitable for downstream clustering and predictive modeling.

### 4.4 Unsupervised Clustering of Tissue Fractions

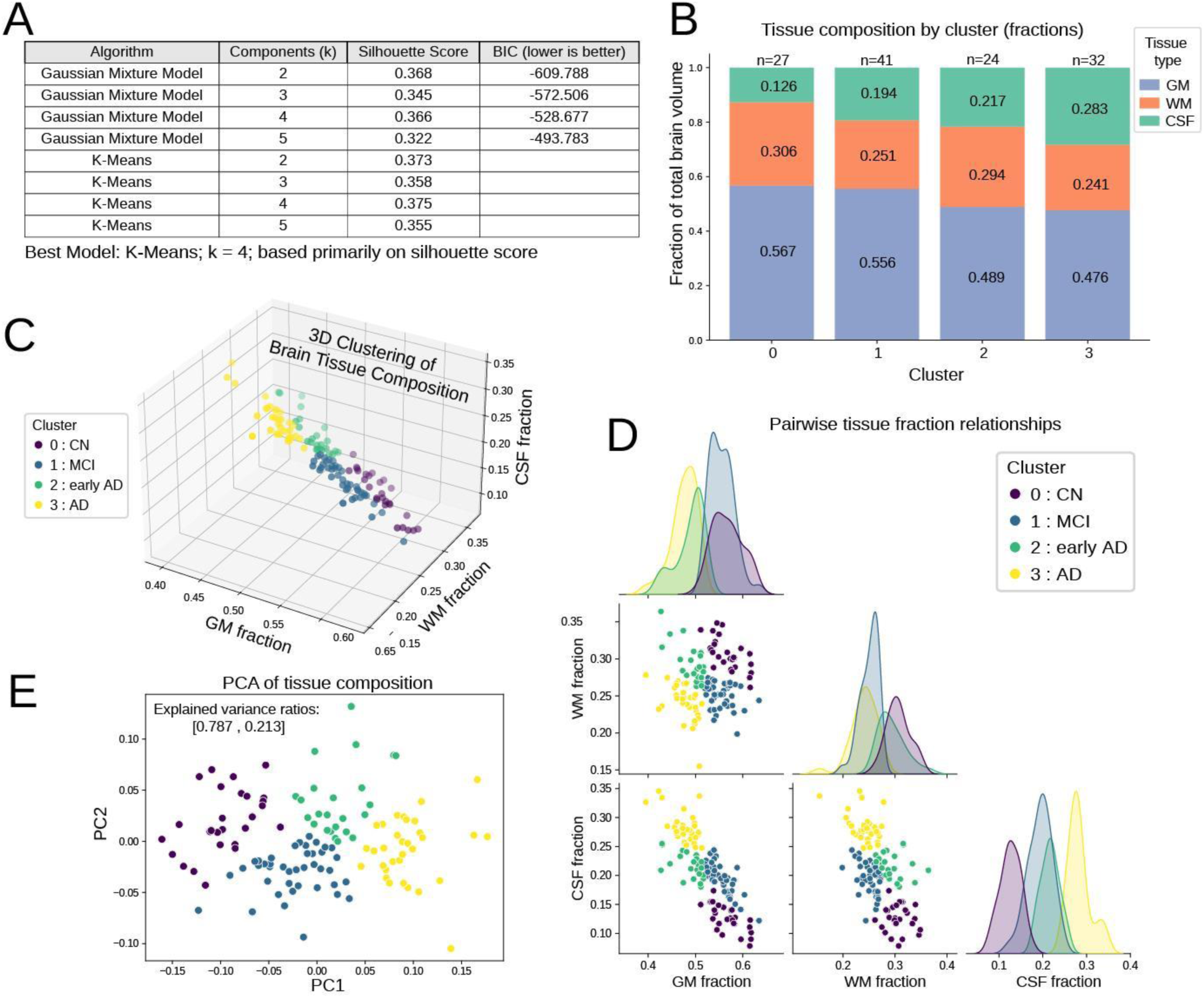

To identify latent neuroanatomical subgroups based solely on tissue composition, an unsupervised clustering analysis was applied to the normalized GM, WM, and CSF fractions. Multiple K-means and GMM solutions were evaluated across cluster numbers *k = 2–5* using silhouette and BIC metrics (Fig. 4A). The highest silhouette score was obtained for K-means with k = 4, indicating an optimal four-cluster structure. Mean tissue fractions within each cluster (Fig. 4B) revealed a clear monotonic trend: GM and WM progressively declined, whereas CSF increased from Cluster 0 → 3, representing a continuum from preserved to atrophic morphologies. This gradient mirrors the biological trajectory from cognitively normal (CN) to mild cognitive impairment (MCI) and Alzheimer’s disease (AD) stages—where GM loss signifies cortical atrophy, WM reduction indicates declining structural connectivity, and CSF expansion reflects ventricular enlargement. The resulting clusters exhibited distinct tissue-composition profiles:

- **Cluster 0** — *Cognitively Normal (CN)-like*: highest GM (≈ 0.57), moderate WM (≈ 0.31), and lowest CSF (≈ 0.13); represents structurally preserved brains.
- **Cluster 1** — *Mild Cognitive Impairment (MCI)-like*: slightly reduced GM (≈ 0.56) and WM (≈ 0.25) with mild CSF increase (≈ 0.19); early signs of parenchymal loss.
- **Cluster 2** — *Early Alzheimer’s Disease (AD)-like*: more pronounced GM loss (≈ 0.49) and WM decline (≈ 0.29), accompanied by CSF expansion (≈ 0.22).
- **Cluster 3** — *Advanced AD-like*: lowest GM (≈ 0.48) and WM (≈ 0.24) with the largest CSF fraction (≈ 0.28); indicative of extensive cortical atrophy and ventricular enlargement.

The 3-D scatter visualization of tissue fractions (Fig. 4.4C) shows distinct separation of clusters along the GM–CSF axis, highlighting that CSF expansion is the dominant discriminant of disease severity. Pairwise fraction plots (Fig. 4.4D) further confirm these relationships: GM and WM are positively correlated, whereas CSF is negatively correlated with both, consistent with compensatory ventricular expansion accompanying cortical and white-matter loss. A two-dimensional PCA projection (Fig. 4.4E) revealed that the first principal component explained > 70 % of variance, capturing the GM-to-CSF trade-off as the primary axis of structural degeneration. Collectively, these unsupervised results demonstrate that tissue-fraction data alone can spontaneously organize into four biologically interpretable morphometric subtypes, forming a continuum from CN → MCI → early AD → advanced AD that closely mirrors known Alzheimer’s disease progression.

### 4.5 Machine Learning Modeling and Framework

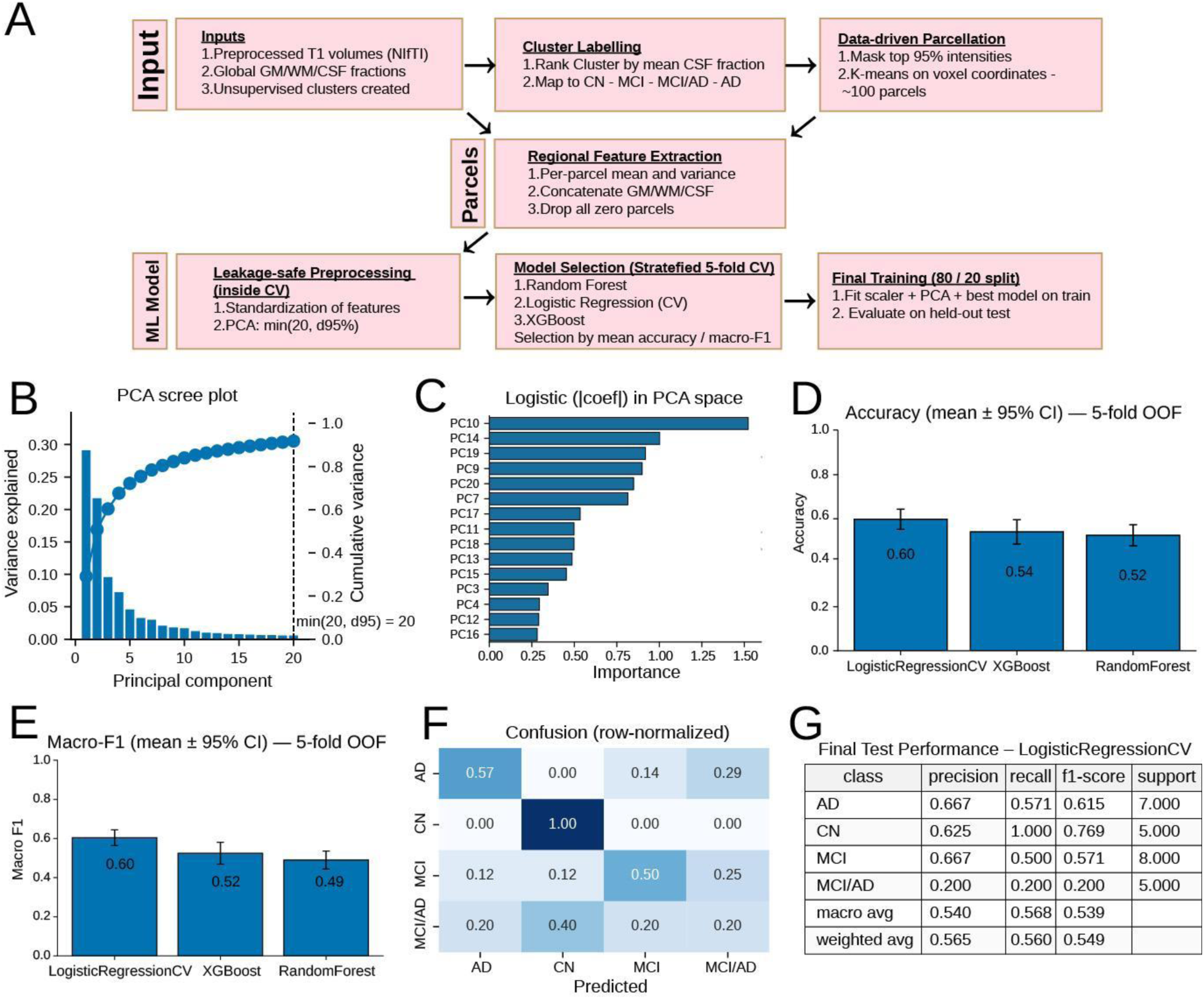

The supervised machine-learning framework built within the MORPH2DIAG pipeline provided a reproducible foundation for classifying morphometric patterns across the Alzheimer’s disease continuum. Starting from relabeled diagnostic groups derived from unsupervised clustering, the workflow as shown in Fig. 5A progressed through data-driven parcellation, extraction of parcel-wise mean and variance intensities, and leakage-safe preprocessing incorporating standardization and PCA embedding. The PCA decomposition in Fig. 5B showed that approximately 95 % of variance was captured within the first 15–20 components, confirming that dimensionality reduction preserved the essential anatomical variability while minimizing noise and redundancy. When visualized in PCA space (Fig. 5C), logistic regression decision boundaries revealed clearly separable clusters for cognitively normal (CN) subjects, while MCI and AD groups exhibited partial overlap, and the MCI/AD subgroup occupied an intermediate zone consistent with its transitional phenotype. Across five-fold cross-validation, Logistic Regression CV achieved the highest generalization accuracy (≈ 0.60 ± 0.03) and macro-F1 (≈ 0.60 ± 0.04), followed by XGBoost (0.54 / 0.52) and Random Forest (0.52 / 0.49) (Fig. 5D–E). This performance hierarchy suggests that linear, regularized models generalized more robustly than tree-based ensembles in the presence of highly correlated morphometric features. The row-normalized confusion matrix in Fig. 5F indicated near-perfect CN identification (∼1.0 recall), moderate separability for AD and MCI (∼0.5 each), and substantial misclassification for the mixed MCI/AD class (∼0.2 correct), reflecting its anatomical and clinical heterogeneity. The final held-out evaluation of the best model, LogRegCV (Fig. 5G), reproduced these trends with stable F1-scores, confirming that CN patterns were consistently the most distinct, while MCI/AD remained the most challenging to distinguish.

### 4.6 Deep Learning Architecture and Framework

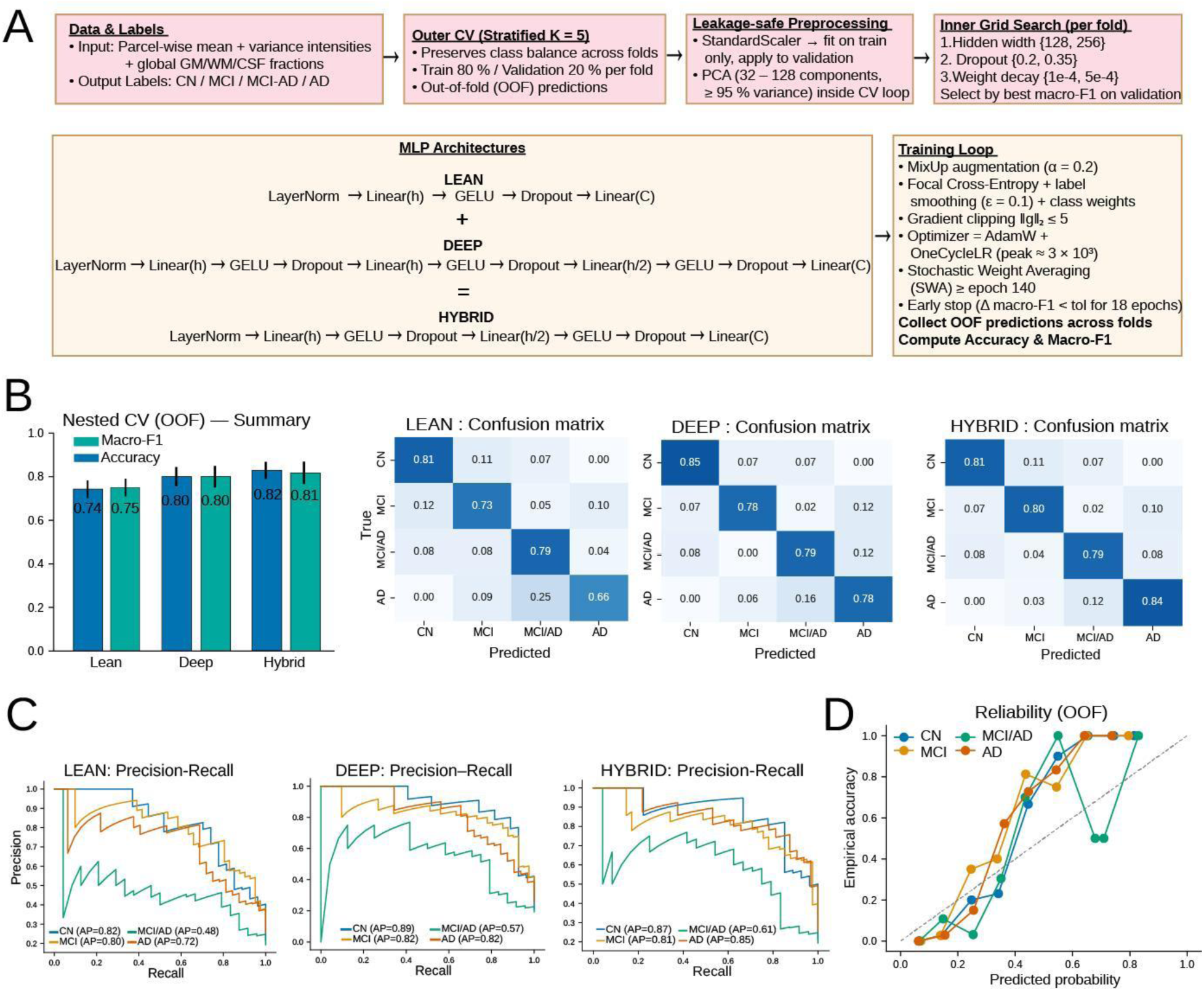

Fig. 6A illustrates the deep-learning stage of the MORPH2DIAG pipeline, spanning data preprocessing, feature representation, model architecture, and evaluation. Input feature matrices comprised parcel-wise mean and variance intensities concatenated with global gray-matter (GM), white-matter (WM), and cerebrospinal-fluid (CSF) fractions. Data were standardized within each training fold, followed by optional PCA reduction (*n* = 32–128 components). Three multilayer perceptron (MLP) variants were trained within a nested cross-validation design:

- Lean MLP: one hidden block (LayerNorm → Linear → GELU → Dropout → Linear).
- Deep MLP: three stacked hidden blocks modeling higher-order nonlinear dependencies.
- Hybrid MLP: intermediate configuration combining the representational width of the deep model with the regularization of the lean network.

Training employed focal-label-smoothed loss with class weighting, AdamW optimization under OneCycleLR scheduling, gradient clipping (‖g‖₂ ≤ 5), and stochastic weight averaging. Early stopping was triggered after 18 epochs without ≥ 1 × 10⁻⁴ improvement in validation macro-F1.

Fig. 6B summarizes the aggregate accuracy and macro-F1 scores across five outer cross-validation folds.

- The Lean MLP achieved ≈ 0.74 accuracy and 0.75 macro-F1,
- the Deep MLP improved to ≈ 0.80 for both metrics,
- and the Hybrid MLP reached ≈ 0.82 accuracy and 0.81 macro-F1, representing the best generalization among all architectures.

Confusion matrices revealed a progressive enhancement in discriminability from Lean → Deep → Hybrid. Deep models reduced cross-class confusion particularly between MCI and MCI/AD, while Hybrid models further increased AD classification performance (> 0.8 true-positive rate). Although CN accuracy slightly decreased relative to Deep, it remained > 0.8, indicating more balanced sensitivity across the disease continuum. Relative to the earlier machine-learning classifiers (macro-F1 ≈ 0.5 - 0.6), the deep models markedly improved separation of intermediate phenotypes—especially MCI and MCI/AD—reflecting higher sensitivity to subtle morphometric variations.

Fig. 6C presents precision–recall curves for each model, showing consistent improvement in both precision and recall as network capacity increased. The Hybrid MLP exhibits steeper curves and larger area under the precision–recall envelope, confirming more reliable detection of minority diagnostic classes and better calibration of probabilistic outputs. Fig. 6D depicts reliability (calibration) plots for the Hybrid model, where empirical accuracy is plotted against predicted probability for each diagnostic label using out-of-fold predictions. Points close to the identity line indicate well-calibrated confidence estimates—predicted probabilities that correspond accurately to observed correctness. Calibration results demonstrate that the hybrid architecture not only improves classification performance but also yields more trustworthy probabilistic predictions—a crucial property for clinical decision support.

## 5. Discussion

This study presents MORPH2DIAG, an automated, atlas-free morphometric pipeline that integrates classical and deep learning approaches to decode structural MRI signatures of neurodegeneration. By standardizing preprocessing, segmentation, and feature extraction, the framework minimizes inter-subject and scanner variability while preserving biologically meaningful morphometric variance^2^. The results demonstrate that structural MRI data alone—when processed through a reproducible, data-driven pipeline—can yield interpretable markers of Alzheimer’s disease progression and cognitive decline^17^.

Machine-learning baselines, including Random Forests, Logistic Regression, and XGBoost, achieved moderate performance (macro-F1 ≈ 0.5–0.6), effectively distinguishing cognitively normal (CN) from impaired individuals but exhibiting consistent confusion between intermediate diagnostic stages (MCI and MCI/AD). This misclassification reflects the biological ambiguity of prodromal Alzheimer’s stages, where regional atrophy and ventricular expansion evolve heterogeneously and are not linearly separable. These findings highlight the limitations of shallow or linear classifiers when confronted with distributed and nonlinear patterns of brain degeneration. In contrast, the deep-learning models showed markedly enhanced discriminative power, achieving up to ∼0.8 macro-F1 accuracy. This improvement underscores the capacity of hierarchical nonlinear transformations to capture spatially distributed morphometric dependencies that conventional models fail to represent. The hybrid multilayer perceptron (MLP) emerged as the most balanced configuration—combining the stability of lean architectures with the representational depth of larger networks—suggesting that moderate model capacity, coupled with robust regularization, yields optimal generalization for structural MRI data. The enhanced sensitivity of the deep and hybrid networks to mild and transitional stages indicates that such architectures are particularly adept at detecting subtle, spatially coordinated atrophy patterns that precede clinically overt Alzheimer’s disease.

Biologically, the hierarchy of model performance (CN > AD > MCI/AD > MCI) mirrors the expected gradation of structural alteration across the disease continuum^17^. CN brains displayed stable cortical and subcortical volumes, while AD subjects showed the most distinct morphometric profiles, marked by pronounced gray-matter reduction and CSF expansion—especially in temporo-limbic and parietal regions.^14^ The deep-learning models were able to recover these spatially distributed signatures, effectively translating them into higher discriminative confidence. The improved classification of AD subjects suggests that the network learned characteristic volumetric and tissue-compositional changes known to accompany advanced disease, while its misclassification of intermediate cases points to overlapping morphometric trajectories during early conversion. From a translational standpoint, these results emphasize the potential of morphometry-based deep learning as a sensitive marker of early neurodegeneration. The ability to identify subtle, nonlinear structural differences suggests that deep feature learning may complement clinical diagnosis by uncovering latent anatomical representations predictive of disease progression. Furthermore, the reproducible, data-driven nature of MORPH2DIAG ensures that such findings remain interpretable and extendable to other datasets or modalities. Collectively, these findings demonstrate that deep-learning approaches—when grounded in standardized, biologically faithful preprocessing—can bridge the gap between descriptive morphometry and predictive neuro diagnostics.

Although MORPH2DIAG derives its morphometric subtypes through unsupervised clustering and later uses these data-driven phenotypes in supervised prediction, this strategy reflects a principled separation between structure discovery and structure validation. The clustering stage isolates latent anatomical patterns inherent to the MRI data—without relying on clinical labels—and parallels recent data-driven approaches that identify biologically coherent axes of neurodegeneration. These subtypes capture reproducible gradients of tissue loss that align with established Alzheimer’s trajectories, providing an anatomically grounded scaffold for subsequent modeling. The supervised and deep-learning components do not replicate the clustering process; instead, they test whether regional and global morphometric features reliably predict these emergent phenotypes in unseen individuals. This division of roles prevents circularity, preserves interpretability, and demonstrates that the discovered structural organization possesses predictive value beyond the initial clustering step.

Importantly, the ability to learn biologically coherent subtypes without requiring diagnostic labels provides a major advantage in the study of neurodegeneration. Early or transitional disease stages—especially prodromal MCI, mixed presentations, and atypical progressors—often exhibit subtle, spatially heterogeneous tissue changes that may not align cleanly with clinical categories. These individuals can be clinically ambiguous yet structurally distinct, and traditional diagnostic labels frequently fail to capture this intermediate biology^19^. By allowing the data to define its own organizational structure, unsupervised morphometric subtyping can reveal patterns of cortical thinning, ventricular expansion, or regional vulnerability that precede overt symptoms, deviate from canonical trajectories, or reflect alternative progression pathways^17^. Such label-free discovery helps identify latent subgroups that may correspond to different disease mechanisms, prognostic risks, or treatment-relevant phenotypes—insights that are rarely accessible from clinician-assigned diagnoses alone^20^.

Thus, MORPH2DIAG not only reduces reliance on imperfect ground-truth labels but also offers a unified and transparent framework that links intrinsic structural variation to clinically meaningful stages of neurodegeneration. This integration bridges descriptive morphometry with predictive neurodiagnostics, positioning the pipeline as a reproducible foundation for future work in population-level subtyping, early detection, and multimodal biomarker development.

## 6. Data & Code Availability

All code and data are available at https://github.com/ShreyaBangera30/MORPH2DIAG. This repository contains all the data and scripts for generating the results.

## Acknowledgements

The authors wish to acknowledge the Department of Statistics, University of California, Berkeley, for its academic support and provision of resources that facilitated this research; and to Dr. Libor Pospisil and Dr. Thomas Bengtsson for their guidance and support throughout the study. The authors also acknowledge Dr. William Jagust and the Jagust Lab at the University of California, Berkeley for their scientific guidance and collaboration; and for facilitating the integration of publicly available neuroimaging datasets into this study.

